# TCR affinity controls the dynamics but not the functional specification of the Th1 response to mycobacteria

**DOI:** 10.1101/2020.10.25.353763

**Authors:** Nayan D Bhattacharyya, Claudio Counoupas, Lina Daniel, Guoliang Zhang, Stuart J Cook, Taylor A Cootes, Sebastian A Stifter, David G Bowen, James A Triccas, Patrick Bertolino, Warwick J Britton, Carl G Feng

## Abstract

The quality of T cell responses depends on the lymphocytes’ ability to undergo clonal expansion, acquire effector functions and traffic to the site of infection. Although TCR signal strength is thought to dominantly shape the T cell response, by using TCR transgenic CD4^+^ T cells with different pMHC binding affinity, we reveal that TCR affinity does not control Th1 effector function acquisition nor the functional output of individual effectors following mycobacterial infection. Rather, TCR affinity calibrates the rate of cell division to synchronize the distinct processes of T cell proliferation, differentiation and trafficking. By timing cell division-dependent IL-12R expression, TCR affinity controls when T cells become receptive to Th1-imprinting IL-12 signals, determining the emergence and magnitude of the Th1 effector pool. These findings reveal a distinct yet cooperative role for IL-12 and TCR signalling in Th1 differentiation and suggests that the temporal activation of clones with different TCR affinity is a major strategy to coordinate immune surveillance against persistent pathogens.

## Introduction

The successful containment of invading pathogens requires the rapid generation of large numbers of antigen-specific T cells with the correct effector function. This involves the activation of distinct programs, including T cell proliferation, differentiation, and migration. Failure in activating or regulating these programs results in impaired host defense. The majority of studies have focused on one or a limited number of CD4^+^ T cell programs, such as, the magnitude of population expansion or expression of master regulators of transcription. There is little information available as to whether these processes, which operate at different biological scales spanning from the molecule to the tissue, are individually or cooperatively regulated *in vivo*. Indeed, *in vitro* studies have already suggested a link between cell division and differentiation, demonstrating that cell division progression is associated with increased expression of signature Th cytokines (1–3).

The pool of naive T cells *in vivo* is diverse and contains clones that express distinct TCRs recognizing different peptide:MHC (pMHC) complexes. It is estimated that there are anywhere between twenty and one thousand naive T cells that possess the same pMHC specificity (4, 5), each with different binding affinities. The strength of TCR signals, regulated by the TCRs affinity, the density of pMHC and co-stimulatory molecules on antigen presenting cells (APCs), regulates downstream T cell activation and function (6, 7). While high affinity TCR signals in cytotoxic CD8^+^ T lymphocytes accelerate cell division and prolong population expansion (8, 9), it delays their migration from secondary lymphoid organs (SLOs) (9, 10) resulting in impaired pathogen control (10). Similarly, strong TCR signals enhance the expansion of CD4^+^ T cell populations (11–13).

Defining the role of TCR signaling strength in the CD4^+^ lymphocyte response is challenging because of the functional heterogeneity in helper T cell populations. The effector function of Th populations is instructed by signals from the TCR as well as from pathogen-conditioned accessory cells and APCs. Historically, investigations into Th cell differentiation have focused on “qualitative” T cell-extrinsic cytokine signals (7, 14). In the case of Th1 differentiation, the innate cytokine IL-12 promotes the generation of interferon-γ (IFN-γ)-producing effectors (15, 16) and host survival following infection with intracellular pathogens (17). Recent studies have suggested a role for “quantitative” differences in TCR signal strength in regulating CD4^+^ T cell differentiation (12, 18–20). Potent TCR signaling is associated with the generation of Th1 (18, 19) or Tfh cells (11, 20, 21). Mechanisms proposed to mediate strong TCR signal-driven Th lineage commitment vary depending on experimental settings. For example, IL-2 (12, 13, 19) and IL-12 receptor signaling (18) have each been suggested to contribute to the generation of Th1 populations following potent TCR stimulation. The model-dependent function of strong TCR signaling suggests a potential interplay between quantitative TCR and qualitative environmental signals in instructing Th differentiation. Currently, the relative role of TCR and innate cytokine signals in lineage commitment and the mechanisms integrating these signals is unknown.

In this study, we developed a T cell adoptive transfer model using CD4^+^ T cells from two TCR transgenic (Tg) mouse lines that recognize the same epitope of the *Mycobacterium tuberculosis* protein Early Secretory Antigenic Target 6 (ESAT-6, E6), with different binding affinities. Following T cell transfer, WT or IL-12-deficient recipient mice were infected with a recombinant *Mycobacterium bovis* Bacillus Calmette-Guérin-expressing E6 (BCG-E6). By tracking transgenic CD4^+^ T cells across multiple time-points and in different tissues following intravenous (i.v.) BCG infection, we reveal that by adjusting the rate of cell division, a major function of TCR affinity is to determine the speed and magnitude of the CD4^+^ T cell response. Moreover, TCR affinity plays a minimal role in specifying T helper cell effector function.

However, by regulating cell division-dependent IL-12Rβ2 expression, TCR affinity controls when T cells become receptive to IL-12 and acquire Th1 effector function. Since high affinity CD4^+^ T cells also migrate to infected non-lymphoid tissues faster than their low affinity counterparts, our findings show that TCR affinity coordinates multiple programs to determine the overall potency of the Th cell response to infection. They also suggest that the temporal activation of distinct T cell clones is a mechanism controlling the initiation and maintenance of Th1 immunity against persistent infection.

## Results

### CD4^+^ T cells with high affinity TCRs are primed earlier and expand further than low affinity cells during mycobacterial infection

To allow a direct comparison of the current and previous studies, which were focused primarily on T cell responses in the spleen following i.v. infection, we developed an i.v. BCG infection and CD4^+^ T cell transfer model (Fig. 1A). BCG delivered by the i.v. route infects primarily the liver and spleen, with the liver being the main site of infection (Fig. S1A). To study the contribution of intrinsic variations in TCR signal strength to the CD4^+^ T cell response, recipient mice were adoptively transferred with GFP-expressing CD4^+^ T cells isolated from two strains of TCR Tg mice (C7 and C24). The TCR of C7 and C24 cells recognize the ESAT6_1-20_ (E6) epitope with low/intermediate and high binding affinity, respectively (22). Recipient mice were then infected with a recombinant BCG that expresses the E6 epitope (BCG-E6) recognized by the T cells. The donor E6-specific CD4^+^ T cells were distinguished from endogenous T lymphocytes by their expression of GFP (Fig. 1B).

**Figure. 1.**
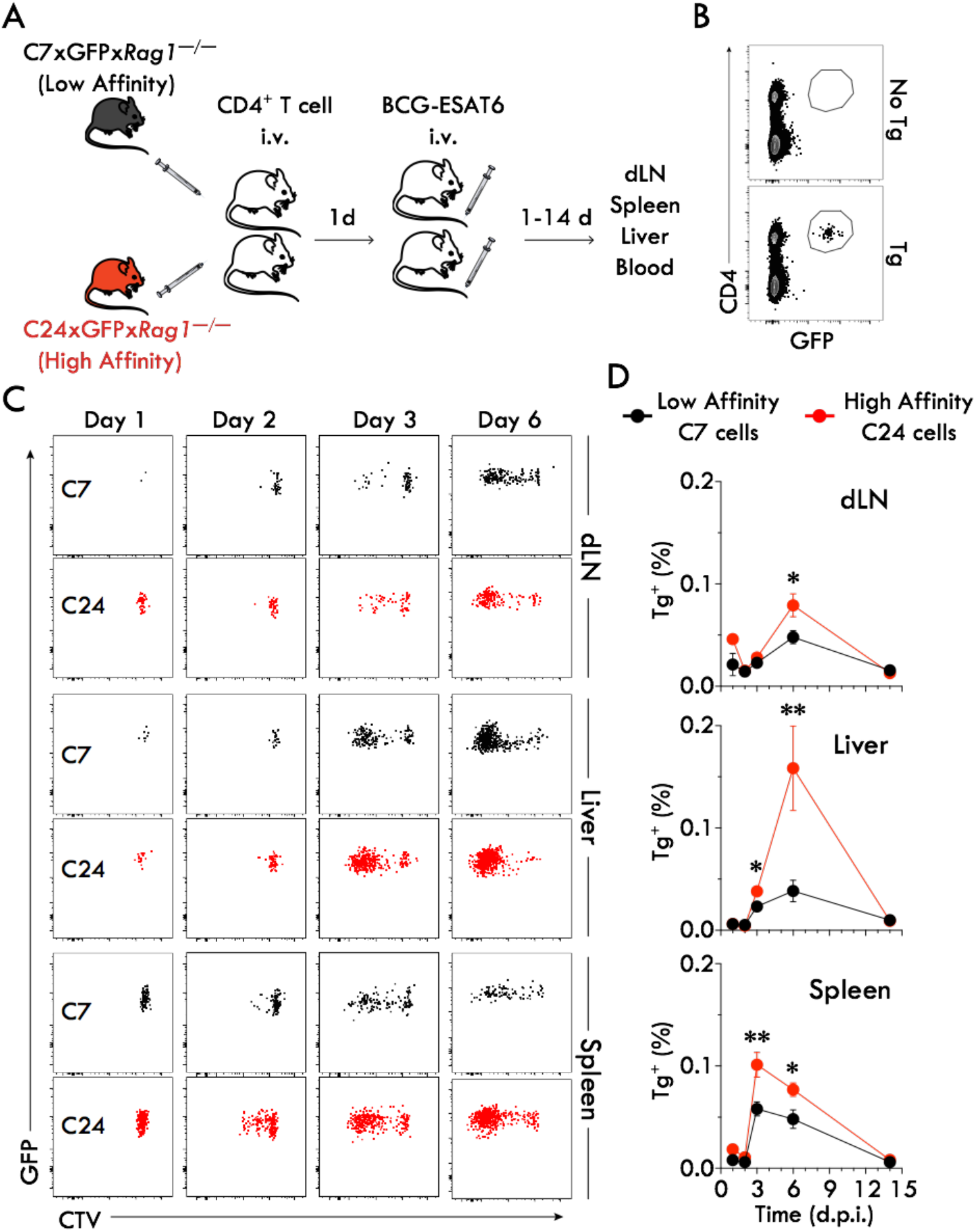
CD4^+^ T cells with high affinity TCRs undergo enhanced accumulation in lymphoid and non-lymphoid organs following mycobacterial infection. **(A)** Schematic diagram illustrating the adoptive transfer and infection model. Magnetically-purified CD4^+^ T cells from C7xGFPx*Rag1*^−/−^ or C24xGFPx*Rag1*^−/−^ TCR transgenic (Tg) mice were transferred i.v. into separate intact, naïve C57BL/6 (B6) recipients (10^5^ / mouse) 1 d prior to i.v. inoculation with BCG-E6 (10^6^ CFU / mouse). The liver-draining lymph node (dLN), spleen, liver and spleen was collected at various time-points p.i.. **(B)** Flow cytometric detection of GFP-expressing TCR Tg T cells in the spleens of recipient mice that did not receive or received transgenic T cells 3 days after BCG-E6 infection. **(C)** Flow cytometric analysis of CTV dilution in GFP-expressing CD4^+^ C7 and C24 cells in the dLN, liver and spleen at day 1, 2, 3 and 6 p.i.. Black and red FACS plots represent C7 and C24 CD4^+^ T cells, respectively. **(D)** Frequency of GFP^+^ C7 or C24 cells in the dLN, liver and spleen at the indicated time-points. Data are pooled from three independent experiments with similar trends; Data shown are the mean percentage ± sem (n > 8 mice / group / time point). Symbols or lines in black and red represent C7 and C24 cells, respectively. Statistical differences between C7 and C24 cells were determined by Student’s t-test (*p<0.05, **p < 0.01).

In this model, both C7 (low affinity) and C24 (high affinity) T cells were detected in the spleen, liver and liver draining lymph nodes (dLN) as early as day 1 after i.v. BCG infection (Fig. 1C). Cell division, assessed by CTV dye dilution, was first detected in Tg cells in the spleen, indicating that the organ is the primary site of T cell priming in this model. CD4^+^ T cell expansion peaked at day 3 post-infection (p.i.) in the spleen compared to day 6 p.i. in the liver and its draining LN (Fig. 1D). Importantly, high affinity C24 cell populations expanded more than lower affinity C7 cells in the spleen, dLN, and liver. Although high affinity C24 T cells demonstrated lower TCR expression when compared to lower affinity C7 cells at day 6 p.i. in the spleen (Fig. S1B), the former Tg T cells exhibited unimpaired IFN-γ expression when re-stimulated with E6 peptide or polyclonal anti-CD3 *ex vivo* (Fig. S1C). These data indicate that TCR downregulation on C24 cells minimally affects their ability to recognize cognate antigens.

### High affinity CD4^+^ T cells undergo accelerated activation, proliferation and differentiation

To investigate whether the accelerated and greater expansion of high affinity C24 cells resulted from their enhanced activation and cell division, we first examined the kinetics of T cell activation and proliferation *in vivo* after adoptive T cell transfer and BCG-E6 infection. A greater fraction of C24 cells upregulated the activation markers CD44 and CD25 than lower affinity clones at day 2 in the spleen (Fig. 2A). By day 3, however, high and low affinity T cells expressed similar levels of CD44. Elevated CD25 expression on CD4^+^ T cells was transient, with CD25 expression returning to negligible levels by day 6 p.i., irrespective of TCR affinity.

**Figure. 2.**
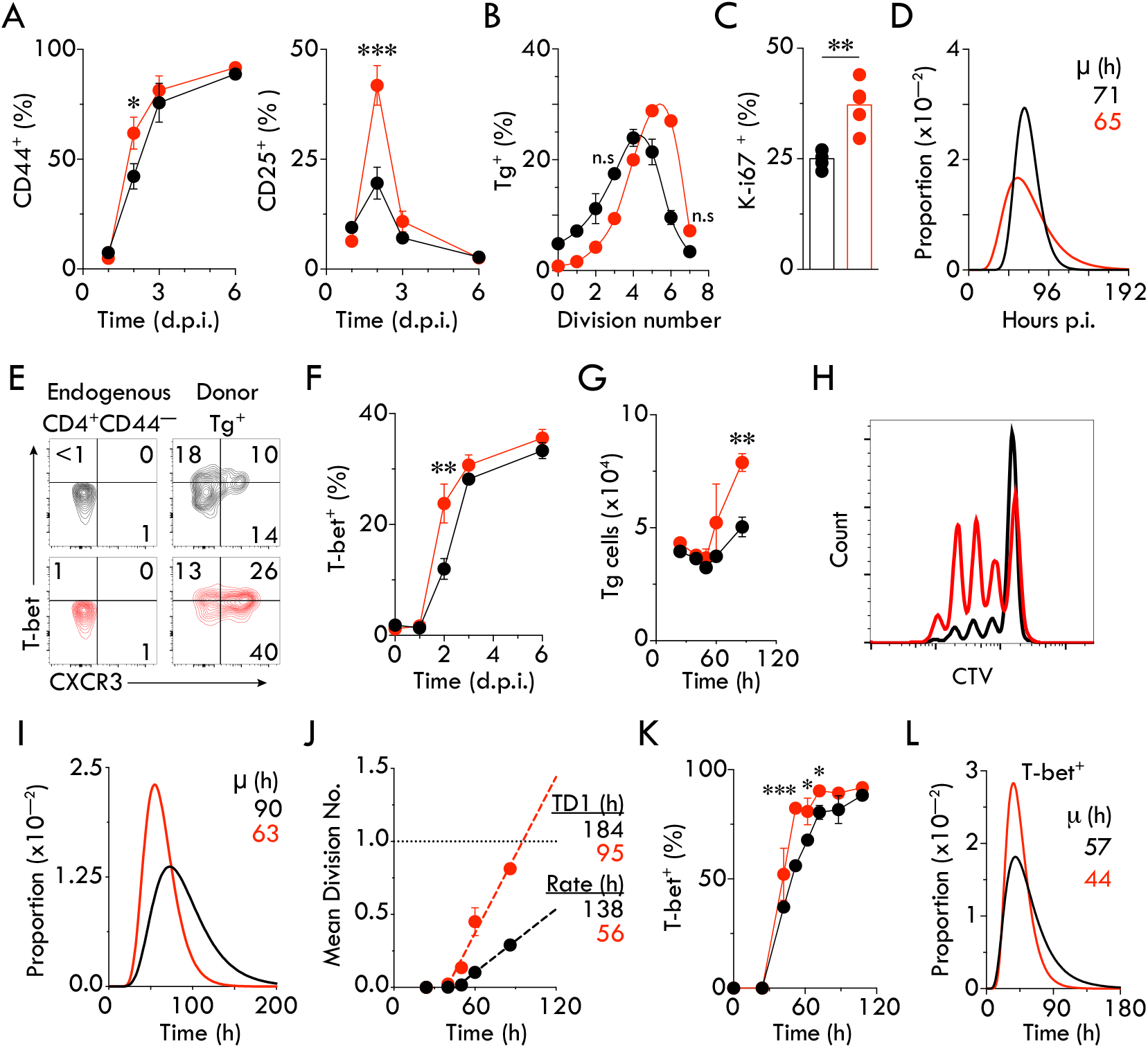
Strong TCR signaling accelerates the activation, proliferation and differentiation of Th1 cells. **(A-F)** CTV-labelled C7 and C24 cells were transferred into naive B6 recipients 1 day prior to inoculation with BCG-E6. The activation and proliferation of TCR Tg cells in the spleen was determined using flow cytometry. **(A)** Flow cytometric analysis of CD44 and CD25 expression on C7 and C24 cells at the indicated time-points. Data are pooled from at least three independent experiments with similar trends. Data shown are the mean percentage ± sem (n > 10 mice / group / time point). **(B)** Frequency of Tg cells in each division 3 d.p.i. determined by flow cytometry. Data are representative of four independent experiments with similar results. Symbols and bars denote the group mean ± sem (n = 3 mice / group), respectively. The difference in the proportion of C7 and C24 cells in each division was statistically significant (at least p<0.05) for all division numbers, unless indicated otherwise. **(C)** Percentage of Ki-67-expressing Tg cells in the spleen 3 days after BCG-E6 inoculation determined by flow cytometry. Closed circles represent individual mice and bars denote the group mean. Data are representative of two independent experiments with similar results (n = 4 mice / group). **(D)** Time to first division determined by CTV dilution *in vivo*. Best-fit log normal probability distribution describing the mean time to first division (C7: μ = 71.1 h, sd. = 0.2 h; C24: μ = 65.2 h, sd. = 0.2 h). Data are representative of two independent experiments (n = 5 mice / group). **(E)** Flow cytometric analysis of T-bet and CXCR3 expression in donor TCR Tg cells (GFP^+^CD4^+^) at 2 d.p.i.. Endogenous naïve CD4^+^ cells (GFP^−^CD4^+^CD44^−^) within the same recipient spleen were used as controls. **(F)** Percentage of T-bet expression at indicated time points p.i.. Data are pooled from three independent experiments Symbols and bars denote the group mean ± sem (3-10 mice / group). **(G-L)** Purified C7 and C24 Tg cells were CTV-labelled and cultured with CD4-depleted splenocytes isolated from naïve B6 mice in the presence of IL-12 (5 ng/mL), anti-IL-4 (10 μg/mL), anti-IFN-γ (10 μg/mL) and the E6 peptide (0.05 μg/mL). **(G)** Accumulation of C7 and C24 Tg cells *in vitro*. Data shown are the mean numbers ± sd of triplicate cultures and representative of three independent experiments. **(H)** Representative flow cytometry plot of CTV dilution in C7 and C24 Tg cells at 60 h. **(I)** Time to first division determined by CTV dilution *in vitro*. Best-fit log normal probability distribution describing the mean time to first division (C7: μ = 89.73 h, sd. = 1.62 h; C24: μ = 62.89 h, sd. = 1.55 h). **(J)** The mean division number (MDN) of precursor C7 and C24 cells at each time point was fitted with a linear regression (C7: y = 0.007224x - 0.3307; C24: y = 0.01784x – 0.6994). The time to first division is calculated by solving for × (time), when the MDN = 1 (C7 = 184.21 h, C24 = 95.26 h). The reciprocal of the slope provides an estimate of the rate of cell division (C7 = 138.43 h, C24 = 56.05 h). **(K)** Kinetic analysis of T-bet expression in C7 and C24 T cells by flow cytometry. Data shown are the mean percentage of T-bet^+^ Tg cells ± sd of triplicate cultures and representative of three independent experiments. **(L)** Quantification of T-bet induction time in Tg cells *in vitro.* Best-fit log normal probability distribution describing the mean time to T-bet expression (C7: μ = 56.5 h, sd. = 1.4 h; C24: μ = 44.2 h, sd. = 1.6 h). Lines and symbols that are black or red represent C7 and C24 cells, respectively. Statistical differences between C7 and C24 cells were determined by Student t-test analysis (*p<0.05, **p < 0.01, ***p < 0.001, ns., not statistically significant).

Consistent with their increased activation, the majority of high affinity C24 cells had undergone at least one additional division when compared to their lower affinity counterparts at day 3 p.i. *in vivo* (Fig. 2B). The enhanced cell cycling of high affinity T cells was further demonstrated by an increased percentage of Ki-67^+^ C24 cells in the spleen (Fig. 2C). Mathematical modelling based on the fraction of divided cells over time (8, 23, 24) (Fig. S2A) estimated that high affinity C24 cells entered cell division 6 h earlier than their lower affinity C7 counterparts *in vivo* (Fig. 2D, 65 h vs. 71 h). One of the hallmarks of the immune response to mycobacterial infection is the generation of Th1 cells. When compared to lower affinity C7 cells, a greater proportion of high affinity C24 cells upregulated T-bet and the Th1-associated chemokine receptor CXCR3 at day 2 p.i. (Fig. 2E). However, these differences were eventually diminished at later time points in the spleen (Fig. 2F). Together with the activation kinetics, these findings reveal that the impact of TCR signal strength on Th1 activation and differentiation is time dependent.

The quantification of T cell proliferation, expansion and differentiation kinetics *in vivo* is confounded by the egress and recirculation of activated lymphocytes. To circumvent these inherent limitations, we developed an *in vitro* system where CTV-labelled Tg T cells were co-cultured with CD4^+^ T cell-depleted WT splenocytes, E6 peptide, and recombinant IL-12. In addition, as IFN-γ enhances the Th1 response *in vitro* (25, 26) but not *in vivo* (27), we included an IFN-γ neutralizing antibody in all the cultures to exclude the role of auto/paracrine IFN-γ in Th1 differentiation.

Recapitulating our findings *in vivo*, high affinity C24 cells underwent greater population expansion (Fig. 2G), as well as accelerated cell division (Fig. 2H) and activation (Fig. S2B) when compared to low affinity C7 cells *in vitro*. At 24 h a greater fraction of high affinity C24 cells expressed molecules associated with early T cell activation, proliferation and metabolic programs, including IRF4 and phosphorylated mTOR (p-mTOR) as well as their downstream target, p-S6 (Fig. S2B). The enhanced activation was also associated with elevated c-myc levels in undivided C24 cells (Fig. S2C). As the extent of c-myc expression in undivided cells can control the number of times a cell divides before it senesces (28), these findings collectively indicate that high affinity TCRs accelerate the activation of proliferative and metabolic programs.

Employing the same mathematical modelling used to determine the time to first division *in vivo*, we estimated that high affinity C24 cells entered their first division approximately 27 h earlier than low affinity C7 cells *in vitro* (63h vs. 90h, Fig. 2I) (Fig. S2D). One caveat associated with this quantification method is that it does not account for how a twofold increase in the number of cells per division would affect the relative ratio of divided and undivided cells over time (8). Therefore, we utilized another well-described method that accounts for the effect of cell division on the total cell number to validate our findings above (8). This method allowed us to track a cohort of cells through different divisions and time-points (Fig. S2E). By calculating the time when the mean division number of a population of precursors is equivalent to 1, the modelling revealed that higher affinity C24 cells took 95 h to enter their first division compared to 184 h for C7 cells (Fig. 2J). Additionally, high affinity TCR signaling reduced the time it took to complete successive divisions (56 h vs. 138 h) (Fig. 2I). Therefore, in addition to reducing the time to first division, our data revealed that high TCR affinity also accelerates the rate of proliferation to enhance the expansion of CD4^+^ T cell populations. Similar to our *in vivo* findings, high affinity C24 populations acquired T-bet earlier than their lower affinity counterparts *in vitro* (Fig. 2K). Mathematical modelling estimated that T-bet was upregulated in high affinity C24 populations 13 h earlier than lower affinity C7 cells (Fig. 2L). These data collectively indicate that CD4^+^ T cells with higher affinity TCRs undergo accelerated cell division and Th1 lineage commitment.

### TCR affinity determines the timing of IL-12-dependent Th1 commitment

Multiple cytokines have been shown to contribute to the development of Th1 cells. In this regard, IL-12 is essential for Th1-dependent immunity against mycobacterial infection in both mice (29) and humans (30). Previous work has suggested that TCR signal strength also plays a role in the acquisition of the Th1 phenotype (12, 19). To examine the relative contribution of TCR versus cytokine signaling in Th1 differentiation, we first compared the kinetics of T-bet induction in high and low affinity CD4^+^ T cells after E6 peptide-stimulation in the presence or absence of exogenous IL-12 *in vitro*. We found that in the absence of IL-12 (in which TCR signaling is the major driver of differentiation), T-bet was only upregulated to an intermediate level in the majority of T cells (Fig. 3A). Kinetic analysis revealed that TCR stimulation alone failed to generate T-bet^high^ (T-bet^hi^) T cell populations regardless of TCR affinity (Fig. 3B). However, the inclusion of exogenous IL-12p70 significantly increased the percentage of T-bet^hi^ populations in both low and high affinity T cell cultures, with the increase accelerated in the latter cultures (Fig. 3A & B). As expected, CXCR3 induction was closely correlated with T-bet expression. When intracellular cytokine expression was evaluated without further re-stimulation, we observed that IFN-γ was predominantly expressed by the cells expressing the highest levels of T-bet, suggesting that T-bet^hi^ cells represent terminally differentiated Th1 effector populations. Importantly, there was an increased accumulation of IFN-γ expressing cells in C24 compared to C7 cultures, suggesting that high TCR affinity accelerates the generation of IL-12-dependent Th1 effectors (Fig. 3C). Kinetic analysis revealed that the IL-12 dependent generation of T-bet^hi^ cells occurred at the expense of T-bet^int^ populations (Fig. S3), irrespective of TCR affinity, such that high affinity TCR signals resulted in the generation of 3 – 4-fold more T-bet^hi^ effectors than their low affinity counterparts (Fig. 3D). Although IL-2 has been shown to enhance Th1 differentiation (31), the inclusion of IL-2 neutralizing antibodies or the addition of exogenous IL-2 did not affect T-bet expression in our system (Fig. S4). These findings suggest that although TCR signals alone can induce T-bet expression, strong TCR signals cannot replace the requirement of IL-12 in the generation of fully differentiated T-bet^hi^ Th1 effectors.

**Figure. 3.**
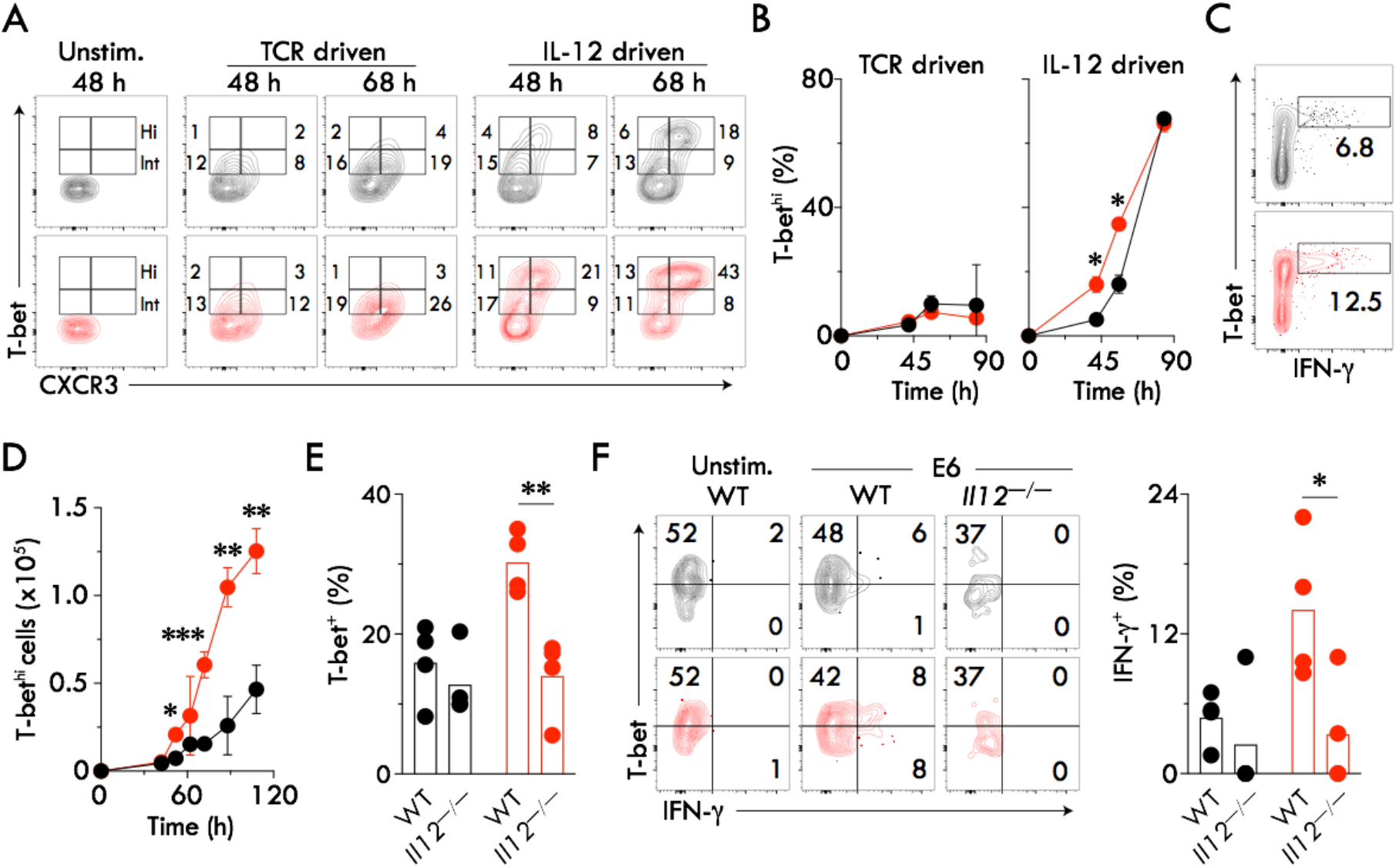
High affinity TCRs accelerate the IL-12-dependent generation of Th1 effector populations. **(A-D)** Purified C7 and C24 CD4^+^ Tg cells were cultured with CD4-depleted splenocytes from naïve B6 mice in the presence or absence of E6 peptide. In some E6-stimulated cultures, mAbs to IL-12p70, IL-4 and IFN-γ were added (TCR-driven). In others, mAbs to IL-4 and IFN-γ, together with recombinant IL-12p70, were included (IL-12-driven). (**A**) Flow cytometry plots showing the levels of T-bet expression in Tg cells in the absence (TCR-driven) or presence of IL-12 (IL-12-driven). (**B**) Changes in the percentage of T-bet high (T-bet^hi^) cells under different polarization conditions analysed by flow cytometry. Data shown are the mean percentage of T-bet^hi^ Tg cells ± sd of triplicate cultures. **(C)** Representative flow cytometry plot showing intracellular IFN-γ expresseion in T-bet^hi^ cell populations. E6 peptide-activated C7 and C24 T cells were cultured under IL-12 driven culture conditions for 66 h and intracellular T-bet and IFN-γ expression was analyzed without further re-stimulation. BFA was added to the culture 6h before analysis. (**D**) Expansion of T-bet^hi^ Tg cell populations in the presence of IL-12, anti-IL-4 and anti-IFN-γ Data shown are the mean numbers of T-bet^hi^ Tg cells ± sd of triplicate cultures. **(E-F)** Purified CD4^+^ T cells from C7xGFPx*Rag1^−/−^* or C24xGFPx*Rag1^−/−^* mice were transferred i.v. into separate naive B6 or *Il12p40^−/−^* recipients 1 day prior to i.v. inoculation with BCG-E6. Data are representative of two independent experiments with similar results (4 mice / group). Symbols denote individual mice and open bars represent group means. **(E)** Proportion of C7 and C24 cells that express T-bet in the spleen at day 2 p.i.. **(F)** Representative flow cytometry plots and summary data showing the percentage of IFN-γ expressing Tg cells in the liver 6 d.p.i.. Hepatic leukocytes were re-stimulated in the presence or absence of E6 peptide (1 μg/mL) for 5 h *ex vivo* before flow cytometry analysis. For all data, the lines, symbols or FACS plots in black and red represent C7 and C24 cells, respectively. Data are representative of three independent experiments with similar results. In **(E)** and **(F)**, closed circles represent individual mice and bars denote the group mean. Statistical differences between C7 and C24 cells were determined by Student t-test analysis, (*p<0.05, **p < 0.01, ***p < 0.001).

To examine whether IL-12 plays a role in determining the Th1 phenotype of low and high affinity CD4 T cells *in vivo*, C7 and C24 cells were adoptively transferred into WT or *Il12p40*-deficient (*Il12^−/−^*) recipient mice, and T-bet expression in donor T cells was assessed following BCG-E6 infection. In contrast to WT recipients, T-bet was not upregulated in the donor TCR Tg T cells in the spleens of *Il12^−/−^* recipient mice regardless of TCR affinity (Fig. 3E). Notably, CD4^+^ T cells with high affinity TCRs failed to display expedited T-bet upregulation in the spleen at day 2 p.i. Similarly, we found that the enhanced ability of the high affinity CD4^+^ T cell populations to express IFN-γ in response to peptide re-stimulation *ex vivo* was compromised if they were transferred into BCG-infected *Il12^−/−^* recipients (Fig. 3F). These findings suggest that IL-12 but not TCR signals are critical for the generation of terminally differentiated Th1 effectors and that the lack of innate signaling cannot be compensated for by strong TCR signals.

### TCR affinity does not regulate the activation status or effector function of individual, divided CD4^+^ T cells

Although TCR affinity temporally regulates the magnitude and function of Th1 populations, it may not regulate the functional output at the level of individual T cells (24, 32, 33). One approach to address this possibility is to compare the mean fluorescence intensity (MFI) of molecules of interest in a given population using flow cytometry (34, 35). However, as low and high affinity T cells are activated with different kinetics (Fig. 2), simply comparing the MFI in bulk populations might skew the analysis in favor of higher affinity populations that have undergone greater division. Hence, we decided to approach this question by comparing the MFI of molecules on Tg cells in the same division.

Activated C7 and C24 T lymphocytes showed comparable expression of CD44, CD25, IRF4 and CXCR3, as well as a similar size when evaluated in the same division *in vitro* (Fig. 4A). This suggests that the activation status of Th cells is not intrinsically regulated by TCR affinity. Similarly, while IL-12 dose-dependent increases in the percentage of T-bet^+^ cells were consistently more predominant in C24 than C7 cultures (Fig. 4B), T-bet levels in the two T cell populations in each cell division were comparable. In agreement with the findings shown above (Fig. 4A and B), enhanced T-bet expression was associated with cell cycle progression and the amount of exogenous IL-12 (Fig. 4C), confirming that TCR affinity controls the dynamics of the Th1 response rather than lineage commitment decisions occurring at the level of individual T cells.

**Figure. 4.**
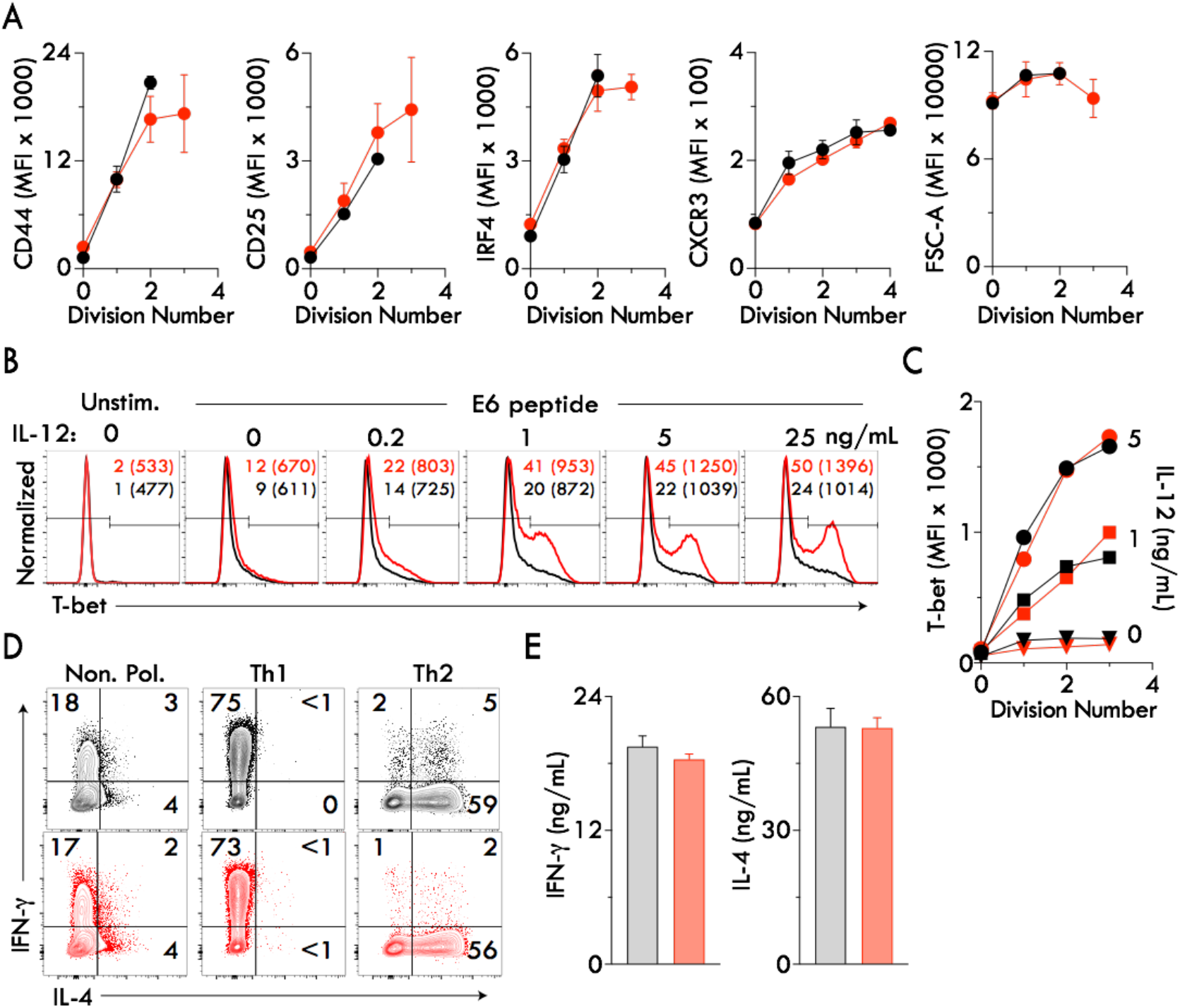
TCR affinity does not determine the activation status or effector function of divided CD4^+^ T cells. **(A-C)** CTV-labelled C7 and C24 Tg cells were stimulated with E6 peptide in the presence of recombinant IL-12p70 and anti-IFN-γ mAb for 72 h. **(A)** Flow cytometric analysis of the MFI of CD44, CD25, IRF4 and CXCR3 as well as FSC-A (cell size) in each division of C7 and C24 populations at 42 h after peptide stimulation. **(B-C)** Tg cells were stimulated with E6 peptide in the presence of the graded concentrations of recombinant IL-12p70 and T-bet expression determined by flow cytometry. **(B)** T-bet expression in C7 and C24 Tg cell populations 60 h after stimulation. Numbers outside and within parentheses represent the percentage of T-bet^+^ cells and the MFI of T-bet expression in T-bet^+^ populations, respectively. (**C**) MFI of T-bet in each division of CTV-labelled C7 and C24 cells 66 h after stimulation. Data shown are representative of two independent experiments with similar results. **(D-E)** Purified C7 and C24 CD4^+^ T cells were stimulated with E6 peptide under non-polarizing (anti-IL-12, anti-IFN-γ and anti-IL-4 mAbs), Th1 (IL-12p70 with anti-IFN-γ and anti-IL-4 mAbs) or Th2 (IL-4 with anti-IFN-γ and anti-IL-12 mAbs) culture conditions for 96 h. Following expansion in IL-2-containing medium for 48 h, an equal number of T cells were re-stimulated with immobilized anti-CD3 in the presence (**D**) or absence (**E**) of BFA for 6 h. **(D)** Intracellular IFN-γ and IL-4 expression under the indicated culture conditions was analyzed by flow cytometry. **(E)** Quantification of IFN-γ and IL-4 produced by Th1 and Th2 Tg cells by Cytokine Bead Array. Data shown are the mean ± the sd of triplicate cultures. For all data the bars, lines, symbols or FACS plots in black and red represent C7 and C24 cells, respectively.

Finally, low and high affinity T cells were equally susceptible to Th1 and Th2 polarization. Following priming with E6 peptide under non-polarizing, Th1 or Th2 culture conditions *in vitro* and re-stimulation with plate-bound anti-CD3 mAb, an equivalent proportion of C7 and C24 cells expressed the expected signature Th cytokines IFN-γ and IL-4 (Fig. 4D). Importantly, when an equal number of cells were re-stimulated, C7 and C24 cell cultures produced comparable amounts of the signature cytokines, indicating that low and high affinity effector T cells have similar cytokine-producing potential at the per-cell level (Fig. 4E). These data indicate that TCR affinity itself does not intrinsically instruct the differentiation of CD4^+^ T cells or determine their activation phenotype.

### TCR signal-controlled IL-12Rβ2 upregulation is cell division-dependent

To determine how IL-12 cooperated with TCR affinity to control the timing of Th1 polarization and ultimately the expansion of effector populations, we next examined whether TCR affinity regulated the expression of the signaling component of the IL-12R, IL-12Rβ2, which is known to be absent on naïve T cells (36). We found that when compared to lower affinity CD4^+^ T cells, a greater fraction of high affinity T cells upregulated IL-12Rβ2 48 h after stimulation (Fig. 5A). While IL-12Rβ2 expression was elevated on both TCR Tg T cells over time, the upregulation was significantly accelerated in high affinity T cells leading to increased accumulation of T-bet^hi^ cells in C24 cultures (Fig. 5B). Importantly, the number of IL-12Rβ2-expressing cells in high affinity C24 cell cultures was consistently more than double of that in C7 cultures after 60 h. Regardless of the strength of TCR signaling, IL-12Rβ2 was predominantly detected in T-bet-expressing cells (Fig 5A).

**Figure. 5.**
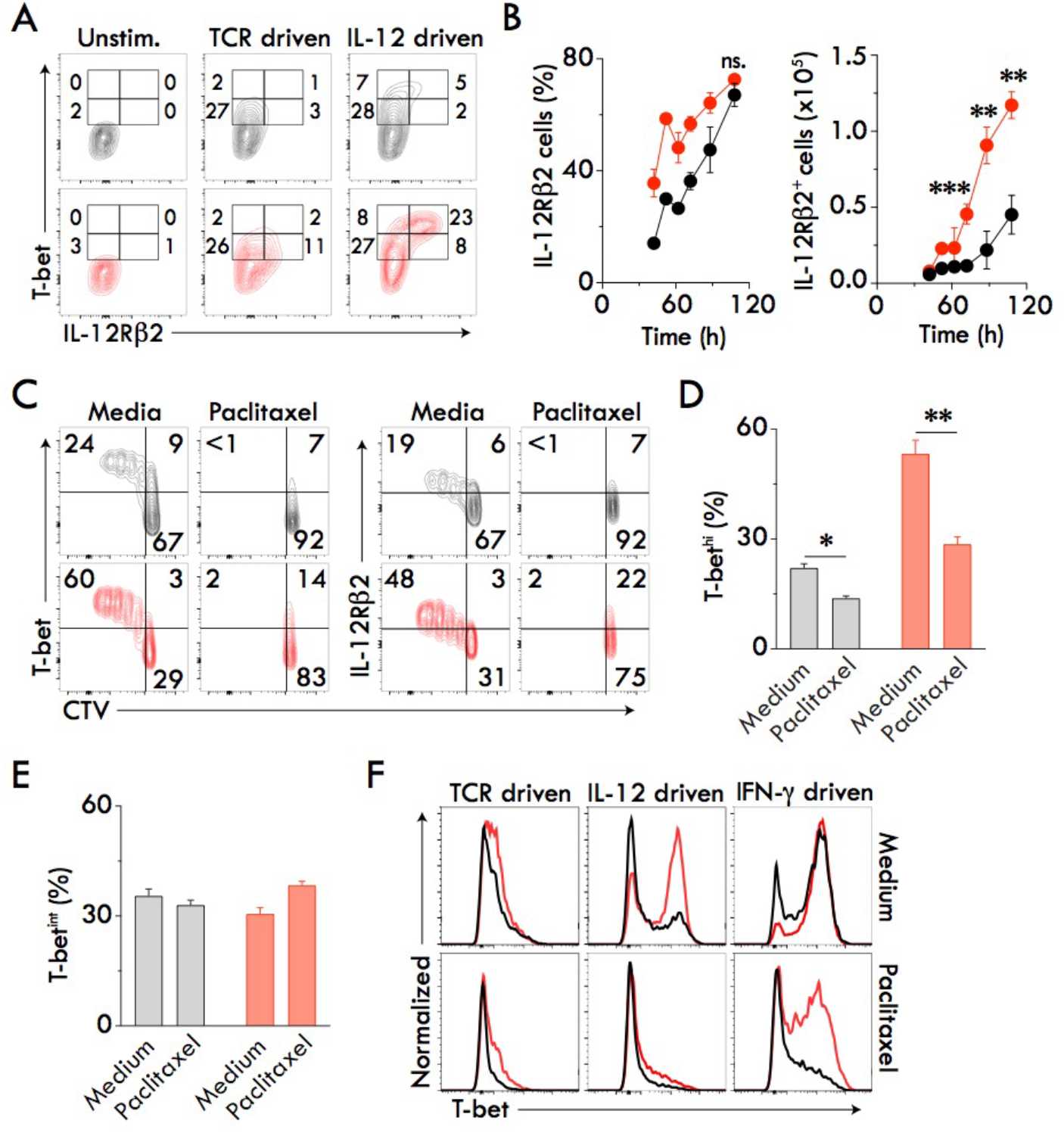
TCR signal-driven IL-12Rβ2 upregulation is cell division-dependent. Purified C7 and C24 CD4^+^ Tg cells were cultured with CD4-depleted splenocytes from naïve B6 mice in the presence or absence of E6 peptide. **(A)** Flow cytometry plots showing IL-12Rβ2 and T-bet expression in Tg cells 48 h after stimulation. In some E6-stimulated cultures, mAbs to IL-12p70, IL-4 and IFN-γ were added (TCR-driven) while in others, mAbs to IL-4 and IFN-γ, together with recombinant IL-12p70, were included (IL-12-driven). **(B)** Kinetic analysis of the percentage and absolute number of IL-12Rβ2-expressing Tg cells by flow cytometry. Data shown are the mean ± sd of triplicate cultures. Statistical differences between C7 and C24 cultures were determined by Students t-test (*p<0.05, **p < 0.01, ***p<0.001, ns., not statistically significant). The difference in the proportion of IL-12Rβ2^+^ C7 and C24 cells was statistically significant (at least p<0.05) for all time points, unless indicated otherwise. **(C-E)** CTV-labelled C7 and C24 T cells were stimulated with E6 peptide and IL-12 in the presence of anti-IL-4 and anti-IFN-γ mAbs. Where indicated, paclitaxel (200 nM) was added at the beginning of the cultures. **(C)** Flow cytometric analysis of T-bet and IL-12Rβ2 expression on CTV-labelled C7 and C24 cells 66 h after peptide stimulation. The percentage of **(D)** Tbet^hi^ and **(E)** T-bet^int^ populations in C7 and C24 cultures with or without paclitaxel examined by flow cytometry at 52 h after peptide stimulation. Bars that are black or red represent C7 and C24 populations, respectively. Data shown are mean ± sd of triplicate cultures. Statistical differences between untreated and paclitaxel-treated cultures were determined by Students t-test (*p<0.05, **p < 0.01). **(F)** Flow cytometric analysis of the effect of paclitaxel treatment on IL-12 versus IFN-γ augmented T-bet expression. As specified, cells were exposed to TCR-driven (anti-IL-12p70, anti-IL-4 and anti-IFN-γ mAbs), IL-12-driven (5 ng/mL IL-12p70 and anti-IFN-γ and anti-IL-4 mAbs) or IFN-γ-driven (anti-IL-12p70, anti-IL-4 mAbs) culture conditions and analyzed at 66 h. All data are representative of at least two independent experiments with similar results.

As the kinetics of T-bet and IL-12Rβ2 induction mirrored that of T cell division (Fig. 2), we hypothesized that TCR affinity timed IL-12Rβ2 expression, and consequent Th1 lineage commitment, by regulating cell division. To test this, Tg CD4^+^ T cells were stimulated with E6 and IL-12p70 in the presence or absence of paclitaxel, a chemical known to inhibit the G2/M phase of the cell cycle (37). As expected, paclitaxel treatment effectively blocked the proliferation of C7 and C24 cells (Fig. 5C). Supporting the key role of cell division in mediating IL-12Rβ2 expression, paclitaxel treatment also impaired the upregulation of IL-12Rβ2 and the subsequent expansion of T-bet-expressing cells. Importantly, paclitaxel treatment was found to selectively impair the generation of T-bet^hi^ cells (Fig. 5D), with T-bet^int^ populations being minimally affected (Fig. 5E). These data place cell division as a mechanism by which TCR affinity fine-tunes the speed of Th1 differentiation. It is worth noting that this cell division-dependent mechanism is not universally utilized in Th cell differentiation. By excluding the anti-IFN-γ monoclonal antibody from the culture, we found that chemical inhibition of cell division prevented IL-12, but not IFN-γ-driven T-bet expression (Fig. 5F). This is likely due to the fact that in contrast to IL-12Rβ2, the IFN-γ receptor is constitutively expressed on naïve T cells (38, 39), and therefore, the effect of IFN-γ on Th1 differentiation is independent of cell division.

### CD4^+^ T cells with high affinity TCRs orchestrate accelerated anti-mycobacterial defense in non-lymphoid tissues

CD8^+^ T cells primed by low affinity antigens have been shown to egress from the spleen earlier than those stimulated by high affinity antigens following *L. monocytogenes* infection (9, 10). However, the role of TCR affinity in coordinating the egress of CD4^+^ T cell egress from secondary lymphoid organs as well as their migration into non-lymphoid tissues is unknown. We first assessed the anatomic localization of Tg CD4^+^ T cells in the spleen of BCG-E6-infected mice on day 2 and 3 p.i.. At day 2 p.i., the majority of T cells, irrespective of TCR affinity, resided mainly within the T cell zone (Fig. 6A), with few GFP^+^ Tg CD4^+^ T cells being detected in the blood (Fig. 6B). By day 3, however, high affinity C24 cells began to egress from the T cell zone, which correlated with their increased presence in B cell follicles and in the blood of recipient mice. In contrast, the majority of lower affinity C7 cells remained in the T cell zone.

**Figure. 6.**
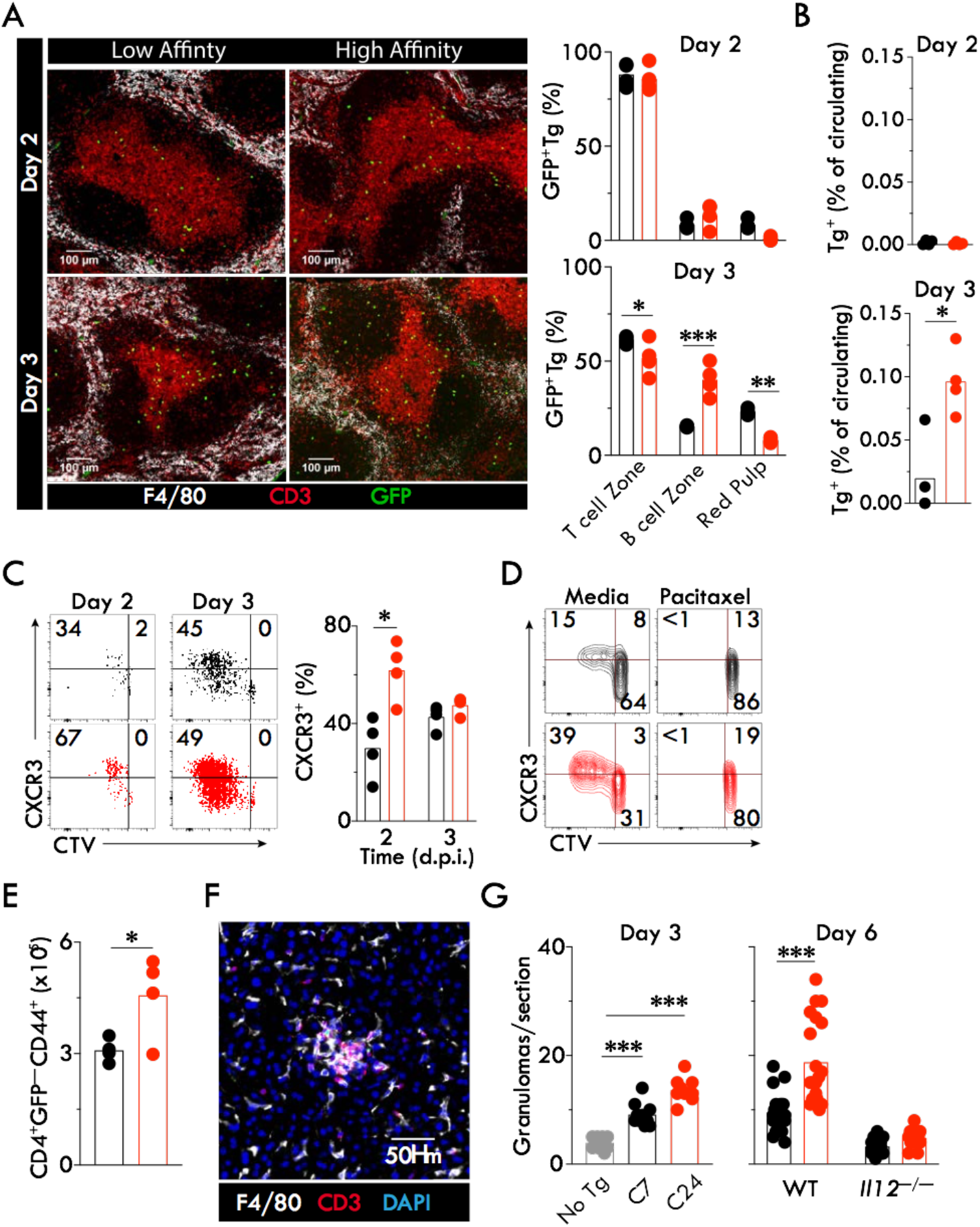
High affinity TCRs accelerate Th1 cell egress from the spleen to expedite the anti-mycobacterial Th1 tissue response at the primary site of infection. CTV-labelled C7 and C24 CD4^+^ cells were transferred into naive B6 recipients 1 day prior to BCG-E6 inoculation. Spleen, blood and liver samples were collected at the indicated time points. **(A)** Microscopic analysis and quantification of the intra-splenic localisation of GFP^+^ Tg cells (green) in the red pulp (F4/80^+^, white), T cell zone (CD3^+^, red) and B cell zone (F4/80^−^ and CD3^−^) at 2 and 3 d.p.i. Histograms depict the number of CD3^+^GFP^+^ cells counted in anatomic site of the spleen. Each symbol represents the percentage of Tg cells per T cell zone, B cell zone or the red pulp of the total number of Tg cells counted in an entire section. Each closed circle denotes one section with bars representing group means. **(B)** Flow cytometric analysis of the frequency of C7 and C24 cells among total CD4^+^ T cell populations in circulation at 2 and 3 d.p.i.. Closed circles denote individual mice and open bars represent group means. **(C)** Flow cytometric analysis of CXCR3 expression on C7 and C24 cells in the spleen at 2 and 3 d.p.i. Data are representative of two independent experiments with similar results (n = 4 mice / group). Closed circles denote individual mice and open bars represent group means. **(D)** Flow cytometric analysis of CXCR3 expression on CTV-labelled C7 and C24 T cells at 66 h after stimulation with E6 peptide, IL-12, and anti-IL-4 and anti-IFN-γ mAbs, in the presence or absence of paclitaxel (200 nM). **(E)** The number of activated endogenous (GFP^−^CD44^+^) CD4^+^ T cells in the livers of WT mice receiving C7 or C24 cells at 6 d.p.i.. Data are representative of two independent experiments with similar results (n = 4 mice / group). **(F)** Fluorescent micrograph showing granulomatous aggregations of F4/80^+^ (white) and CD3^+^ (red) cells in the liver of BCG-E6-infected mice at 3 d.p.i.. **(G)** The enumeration of the total number of hepatic granulomas (CD3^+^DAPI^+^F480^+^ aggregates) in BCG-E6-infected WT or *Il12p40^−/−^* mice. Each closed circle denotes the number of granulomas in an individual section, with twelve sections counted per group. Bars represent group means. Bars, symbols or FACS plots in black and red represent C7 and C24 cells, respectively. Statistical differences between C7 and C24 cells were determined using the **(A, G)** two-way ANOVA or by **(B, C, E)** the students t-test (*p< 0.05, **p<0.01, ***p< 0.001).

T cells traffic from lymphoid tissues into peripherally infected non-lymphoid tissues where they perform their effector function. The chemokine receptor CXCR3 has been shown to be critical for T cell entry into inflamed peripheral tissues (40), and its optimal expression is dependent on T-bet (41, 42). Our data confirmed a strong correlation between the expression of T-bet and CXCR3 (Fig. 2E and 3A). Following the adoptive transfer of CTV-labelled TCR Tg CD4^+^ T cells and i.v. infection with BCG, a greater proportion of CD4^+^ T cells with high affinity TCRs expressed CXCR3 in the spleen when compared to those with lower affinity TCRs at day 2 p.i (Fig. 6C). By day 3 p.i, the proportion of CXCR3^+^ T cells became comparable in C7 and C24 populations, which could be due to the egress of high affinity CXCR3^+^ C24 cells. Enhanced CXCR3 expression was positively correlated with T cell proliferation, which indicates that cells that have undergone multiple rounds of division are better equipped for peripheral tissue entry. To formally establish whether CXCR3 upregulation is dependent on T cell division, we assessed the expression of CXCR3 on peptide-activated, CTV-labelled TCR Tg T cells in the presence or absence of the cell division inhibitor paclitaxel *in vitro*. We confirmed that similar to T-bet, optimal CXCR3 upregulation appeared to be dependent on cell division (Fig. 6D). This suggests that cell division equips CD4^+^ T cells with the chemokine receptor that increases their entry into infected non-lymphoid tissues, with the process temporally regulated by TCR affinity. The accelerated egress of high affinity C24 cells from the spleen was associated with a greater influx of activated CD44^+^ endogenous CD4^+^ T cells into the liver (Fig. 6E), indicating that high affinity CD4^+^ T cells promote the recruitment of endogenous CD4^+^ T cells.

Following mycobacterial infection, a major function of effector Th cells is to organize the formation of granulomas, a key feature of the tissue response involved in containing the infection (43). Hepatic granulomas in BCG-infected mice are characterized as aggregates of F4/80^+^ macrophages and CD3^+^ lymphocytes (Fig. 6F), which gradually mature in size and number following i.v. BCG inoculation (44). While the number of granulomas in the livers of non-T cell transferred mice were low at day 3 p.i., this increased significantly in mice that received TCR Tg CD4^+^ T cells (Fig. 6G). The number of granulomas in the livers of mice receiving C24 cells were significantly higher than those that received low affinity C7 cells, which is consistent with the observation that the egress of high affinity T cells from the spleen was accelerated (Fig. 6B). Finally, we demonstrated that the donor T cell-enhanced formation of hepatic granulomas is dependent on IL-12 regardless of TCR affinity (Fig. 6G). Together, these data reveal that TCR affinity coordinates T cell trafficking and accelerated host defense strategies within infected tissues.

## Discussion

Recent investigations have suggested that TCR signal strength (regulated intrinsically in T cells by TCR affinity, and extrinsically by the density of pMHC and co-stimulatory molecules on APCs) plays a key role in regulating T cell responses to a variety of infections. However, mechanisms underlying the T cell-intrinsic regulation of lymphocyte responses, particularly in CD4^+^ T cells, are incompletely understood. In this regard, whether strong TCR signals could replace the requirement of environmental cues in imprinting the effector function of CD4^+^ T cells was undefined. We reveal that TCR affinity neither quantitatively regulates the functional output nor directly determines Th1 lineage commitment in individual CD4^+^ lymphocytes. Instead, by controlling cell division-dependent IL-12R expression, TCR affinity controls when T cells become sensitive to environmental cues, and tailor their effector function to the microbe encountered. We suggest that the primary function of TCR affinity in the CD4^+^ T cell response to intracellular infection is to control the speed of Th1 differentiation and the magnitude of effector cell populations. Interestingly, a similar observation has been reported with CD8^+^ T cells (32, 33, 45). This cell division-dependent mechanism also helps explain how the diverse processes of a T cell response, proliferation, differentiation and trafficking, are temporally synchronized by TCR signals.

The TCR affinity-regulated, time-dependent mechanism is particularly relevant to the understanding of host-pathogen interactions in diseases caused by persistent intracellular pathogens, such as *M. tuberculosis* and *Leishmania major*. These pathogens are slow-growing and can initiate a chronic infection with very small inoculants at the site of entry (46, 47). As shown by intravital imaging analysis, antigen presentation during mycobacterial infection is limited (44). Hence, CD4^+^ T cells with high TCR affinity will have a significant advantage over their low affinity counterparts in recognizing poorly expressed microbial peptides, and in receiving IL-12 signals to stabilize the effector function of Th1 cells. Although antigen-affinity has previously been linked to the intra-splenic localization of CD4^+^ T cells (12), whether TCR stimulation strength controls Th cell migration to infected peripheral tissues has not been investigated. Our analysis demonstrates that high affinity CD4^+^ T cell populations infiltrate the liver earlier and in greater numbers, leading to an augmented granulomatous response critical for containing mycobacteria. Interestingly, this was associated with the increased recruitment of activated endogenous CD4^+^ T cells, suggesting that an additional function of Th cells with high affinity TCRs is to enhance the recruitment of other pathogen-specific T cells to the site of infection. CD4^+^ T cells with high affinity TCRs likely play a pivotal role in early pathogen containment because of their ability to undergo accelerated priming, clonal expansion, differentiation and trafficking to the site of infection. Although slower at entering cell division, once activated, low affinity CD4^+^ T cells are equipped with effector functions that are indistinguishable from high affinity T cells. Therefore, the sequential activation of CD4^+^ T cells with high and low affinity TCRs will not only provide robust but also sustained immunity against persistent pathogens.

This study establishes that TCR signals alone play a minimal role in instructing Th lineage commitment. This notion is further supported by the observation that peptide-activated low and high affinity CD4^+^ T cells were equally capable of becoming Th2 cells when exogenous IL-4 was present. This finding seems to be unexpected because of the general belief that strong TCR signals promote Th1 differentiation, whereas weak signals favor the generation of Th2 cells (48, 49). It is possible that TCR pMHC binding affinity has a distinct function in Th differentiation compared to other TCR signal strength modulators, such as antigen dose or co-stimulatory molecules, as proposed previously (12). However, the discrepancy in the role of TCR affinity in instructing Th differentiation between the current and previous studies could be reconciled if the temporal regulation of T cell responses by TCR signal strength was taken into account. Since the majority of previous studies have reported T cell responses at the population level, in bulk cultures, at a single time point or tissue, the spatiotemporal difference in the response of low and high affinity T cells may not have been captured. The enhanced Th1 response associated with strong TCR stimulation at a single time point reported previously could be due to the accelerated increase in activated cell populations rather than enhanced lineage specification in individual cells.

In agreement with our findings, a previous study has shown that by modulating antigen dose or costimulatory molecule expression, strong TCR signals enhance the expression of IL-12Rβ2 (18). We identified the upregulation of IL-12Rβ2 was dependent on cell division and timed by TCR signal strength, suggesting that cell division is an essential mechanism integrating intrinsic TCR signals and extrinsic environmental cues during Th1 differentiation. Consistent with a previous study (50), the upregulation of IL-12Rβ2 occurs late after T cell activation and is persistently expressed, suggesting that the cells remain susceptible to IL-12 instruction for a sustained time period. Our findings support a model where TCR signals are a digital ‘switch’ that grant individual Th1 cells the ability to receive the instructive and selective cytokine signals for terminal differentiation and the stability of effector cell populations (51, 52). Indeed, IL-12 is required for the maintenance of immunity against a variety of pathogens (53–56). However, it is worth noting that the cell division-dependent mechanism underlying Th1 differentiation may not be generalized for other Th subsets. For example, IL-4Rα, constitutively expressed by naïve T cells, is transiently upregulated following TCR stimulation (57) but subsequently downregulated (57, 58). Therefore, the link between cell division and cytokine receptor expression may not be necessary for the differentiation of Th2 cells.

The critical requirement for environmental cues in instructing Th1 differentiation also suggests that the conclusions of studies investigating the role of TCR signal strength *in vivo* are likely influenced by the infection model. Th1 differentiation associated with strong TCR stimulation during infection with influenza virus, *L. monocytogenes* or Lymphocytic choriomeningitis virus (LCMV) (12, 13, 19) has been attributed to IL-2/STAT5 signaling (12, 20), due partly to persistent CD25 expression on CD4^+^ T cells. While the role of IL-12 was not investigated in these studies, it is likely these pathogens activate distinct innate and adaptive immune responses compared to mycobacteria. IL-12 is only transiently or minimally produced following *L. monocytogenes* and LCMV infection (59). Moreover, IL-12 is either partially required (*L.monocytogenes* or influenza virus) or dispensable (LCMV) for Th1 differentiation *in vivo* (60–63). Importantly, in contrast to mycobacterial infection, host control of the above-mentioned microbes predominantly requires CD8^+^, rather than CD4^+^ T cells (64–67). Interestingly, CD25 was only transiently upregulated in our model. It is possible that the IL-2/CD25 feed-forward loop required for the maintenance of CD25 expression on CD4^+^ T cells is suppressed by mycobacterium-induced IL-12, as IL-12-dependent STAT4 activation and T-bet expression have been shown to suppress IL-2 production and IL-2R signaling (68, 69).

The findings reported emphasize the critical importance of TCR affinity in the timing and scaling of CD4^+^ T lymphocyte responses. They also clarify the relative roles of TCR and IL-12 signaling in Th1 differentiation. Recognizing cell division as the coordinator of distinct components of the CD4^+^ T cell response unifies the mechanisms that determine potent Th1 immunity. The presence of TCRs with distinct affinities for the same epitope within the immune repertoire ensures that immune surveillance against invading pathogens is rapid, sustained and independent of the level of antigen expression or stage of infection. CD4^+^ T cells with high affinity TCRs may orchestrate local immunity by recruiting and helping other immune populations, such as lower affinity helper and cytotoxic T cells, as well as B lymphocytes. Targeting mechanisms known to increase TCR signal strength, such as increasing co-stimulatory signals or modifying peptide and MHC binding affinity, may lead to the development of more effective vaccination strategies for protection against intracellular pathogens.

## Materials and Methods

### Mice

Clone 7 (C7) and Clone 24 (C24) TCR transgenic (Tg) mice specific for ESAT-6_1-20_ - IA^b^ complexes (kindly provided by Drs Eric Pamer and Michael Glickman) were generated on a C57BL/6 (B6) background as previously published (22). C7 and C24 mice were subsequently bred with GFP-expressing B6 *Rag1*^−/−^ mice to generate C*7.Rag1*^−/−^.GFP and C24. *Rag1*^−/−^.GFP mice, respectively. The TCR transgenic animals and *Il12p40^−/−^* mice were bred and maintained at the Centenary Institute Animal Facility. B6 recipient mice (6-8 weeks old) were purchased from Australian BioResources (Moss Vale, NSW, Australia).

### Mycobacterial growth and infection

*M. bovis* BCG expressing a dominant epitope of the the 6 kDa Early Secretory Antigenic Target (ESAT-6) peptide (ESAT-6_1-20_, peptide sequence QQWNFAGIEAAASA) (termed BCG-E6) was generated using a method described previously (70). The bacterium was grown to log phase at 37°C in Middlebrook 7H9 broth (BD Biosciences, North Ryde, NSW, Australia) supplemented with 10% albumin-dextrose-catalase (ADC), 0.5% glycerol, 0.05% Tween80 and 50 μg/mL of kanamycin and 50 μg/mL hygromycin (Sigma-Aldrich, North Ryde, NSW, Australia). BCG-E6 was washed in PBS and injected intravenously (i.v.) into wild-type B6 or *Il12p40^−/−^* mice (10^6^ colony forming units (CFU) / mouse). BCG numbers in inoculates and tissue homogenates were quantified as CFU using Middlebrook 7H11 agar (BD Biosciences) supplemented with oleic acid-ADC (OADC) and 0.5% glycerol.

### Preparation of single cell suspensions from tissues

Liver-draining lymph nodes and spleens were collected in RPMI 1640 supplemented with 2% FCS (RP2) and dissociated through 70 μm strainers. For leukocyte cell isolations from the liver, euthanized mice were first perfused with 10 mL PBS, and the livers dissociated through a 70μm strainer. After washing with PBS, Liver leukocytes were enriched using 35% Percoll (Cytiva, Marlborough, MA, USA). For blood leukocytes, up to 700 μL of blood was collected via cardiac puncture into EDTA coated collection tubes (Greiner Bio-one, Kremsmünster, Austria). Peripheral blood mononuclear cells (PBMCs) were isolated by gradient centrifugation on Histopaque1083 (Sigma-Aldrich). Erythrocytes in single cell suspensions were lysed with ACK lysis buffer. Cells were washed in RP2 prior to viable cells being counted using trypan blue exclusion on a haemocytometer.

### Isolation, Labelling, Enrichment and Adoptive Transfer of TCR Tg CD4^+^ T cells

Lymph nodes (inguinal, axillary, brachial and mesenteric) and spleens were collected from C7x *Rag1*^−/−^xGFP or C24x *Rag1*^−/−^xGFP mice and macerated through a 70μm strainer to obtain single cell suspensions. Red blood cells were lysed using ACK Lysis Buffer (Thermo Fisher Scientific, North Ryde, NSW, Australia). Where indicated, 2.5×10^7^ cells/mL cells were subsequently labeled with 5 μM Cell Trace Violet (CTV) (Thermo Fisher Scientific) in warm PBS supplemented with 0.1% FCS for 20 minutes in the dark at 37°C. CTV labelling was terminated by adding equal volumes of cold FCS. The cell suspensions were incubated with running buffer (PBS supplemented with 0.5% FCS and 2 mM EDTA) and L3T4 microbeads (Miltenyi Biotec, Maquarie Park, NSW, Australia) according to the manufacturer’s instructions. CD4^+^ T cell fractions were isolated using the AutoMacsPro (Miltenyi Biotec). Enrichment of C7/C24 CD4^+^ T cells was confirmed by flow cytometry with a purity of ~95%. CD4^+^ T cells from female mice were washed in PBS, and 10^5^ i.v. injected into B6 or *Il12p40^−/−^* mice 24 h before BCG-E6 infection.

### *In vitro* T cell culture

CD4^+^ T cells were cultured in RPMI 1640 medium supplemented with 10% FCS, 100 U/ml Penicillin, 100 μg/ml Streptomycin, 25 mM Hepes (Thermo Fisher Scientific) and 50 μM 2ME (RP10). CD4^+^ T cell-depleted B6 splenocytes, prepared using the CD4 Positive Selection kit II (StemCell Technologies, Tullamarine, VIC, Australia), were used as antigen presenting cells. Purified CD4^+^ T cells were added to splenocytes in a ratio of 1:4. Prior to the addition of CD4^+^ T cells, splenocytes were cultured with E6_1-20_ peptide (0.05 μg / mL, GenScript) in round-bottomed 96 well plates (Corning) for 30 minutes at room temperature. Where indicated, IL-12 (0.2-25 ng/mL, Thermo Fisher Scientific), IL-4 (10 ng/mL, PeproTech), human IL-2 (10 IU/mL, PeproTech), Paclitaxel (200 nM, Sigma-Aldrich), rat anti-mouse IFN-γ (10 μg/mL, clone XMG1.2), rat anti-mouse IL-4 (10 μg/mL. clone 11B11), rat anti-mouse IL-12p40/p70 (10 μg/mL, clone C17.8), rat anti-mouse IL-2 (10 μg/mL, clone S4B6) (all from BD Biosciences), were included in the cultures. For analysis of cytokine expression without re-stimulation, 1:1000 Golgi-plug (BD Biosciences) was added to the culture 5 h prior to flow cytometric analysis. For detection of intracellular cytokine expression after re-stimulation *ex vivo*, up to 5×10^6^ leukocytes were incubated with 5 μg/mL E6 peptide or 1 μg/mL α-CD3e (clone 145-2C11, BD Biosciences) in the presence of 1:1000 Golgi-plug for 5 hours. Where indicated, Tg cells were rested for 2 d in recombinant human IL-2 (10 IU/mL) prior to re-stimulation with plate-bound α-CD3 for 6 h in the presence or absence of 1:1000 Golgi-plug. IFN-γ and IL-4 protein in culture supernatants were quantified using the Cytokine Bead Array according to the manufacturer’s instructions (BD Biosciences)

### Flow Cytometry

All samples were acquired on the BD Fortessa using FACSDiva software (BD Biosciences) and analysis was performed using FowJo 10 (BD, Franklin Lakes, NJ, USA). Up to 5×10^6^ cells were washed in cold FACS wash (PBS supplemented with 2% FCS and 2mM EDTA) prior to being stained with a surface receptor antibody cocktail containing FcBlock (BD, 2.4G2) and LIVE/DEAD fixable blue dead cell stain (Thermo Fisher Scientific) for 30 minutes at 4°C. For analysis of IL-12Rβ2 expression, cells were stained separately in FACS wash containing IL-12Rβ2-Biotin (REA200, Miltenyi Biotec) and FcBlock for 30 minutes at 4°C. Where appropriate, cells were stained in FACS wash containing streptavidin PE-Cy/7 conjugates for 20 minutes at 4°C before being incubated with the surface antibody cocktail. The following monoclonal antibodies were used for the detection of cell surface markers: CD4 (RM4-5), CD44 (IM7), CD25 (PC61) (all from BD Biosciences); and CXCR3 (CXCR3-173, BioLegend, San Diego, CA, USA). Cells were washed in FACS buffer prior to acquisition.

For detection of intracellular transcription factors and cytokines, surface stained cells were fixed with 100 μL Cytofix/Cytoperm (BD Biosciences) for 15 minutes at 4°C to preserve GFP expression. To improve transcription factor staining, in some experiments, cells were then incubated with 100 μL 1x Fixation/Permeabilization solution (Thermo Fisher Scientific) for 30 minutes at 4°C. Cells were thoroughly washed in 1x Permeabilization buffer (Thermo Fisher Scientific). Cells were incubated for 1 hour at 4°C in 1x Permeabilization buffer containing a cocktail of the following monoclonal antibodies: IFN-γ (XMG1.2), IL-4 (11B11) (both from BD Biosciences); T-bet (4B10, Biolegend), Ki-67 (SolA15) and IRF4 (3E4) (both from Thermo Fisher Scientific). Cells were washed in 1x Permeabilization buffer and resuspended in FACS buffer prior to acquisition.

For detection of phosphorylated proteins and myc, at the indicated time points, cells were immediately fixed and permeabilized using the Transcription Factor Phospho Buffer set (BD Biosciences) as per the manufacturers instructions. For analysis of myc expression, cells were incubated in 1xPermbuffer III (BD) containing purified anti-myc (clone D84C12, Cell Signaling, Danver, MA, USA) or rabbit IgG isotype matched control for 1 hour at 4°C. Cells were washed and then incubated with polyclonal anti-rabbit IgG (H+L) (Thermo Fisher Scientific) for 1 hour at 4°C. For the detection of phosphorylated proteins, cells were incubated with 1xPermbuffer III containing monoclonal antibodies against phosphorylated-S6-Ribosomal protein (D57.2.2E, Cell Signaling) or phosphorylated-mTOR (clone MRRBY, Thermo Fisher Scientific) for 30 minutes at 4°C. Cells were thoroughly washed in 1x Permbuffer III, FACS wash and then incubated in FACS wash containing monoclonal antibodies targeting cell surface markers and FcBlock for 30 minutes at 4°C. Cells were thoroughly washed in FACS wash prior to acquisition.

### Organ collection and processing for imaging

Spleens and the caudate lobe of livers were collected for microscopic imaging. The spleens were cut into quarters. Tissues were then placed in 5 mL of 4% PFA for 12 hours at 4°C. Fixed tissues were then transferred into 30% (v/v) sucrose (Sigma-Aldrich) in PBS. After 24 hours tissues were embedded in Optimal Cutting Temperature (OCT) compound (VWR Chemicals, Atlanta, GA, USA) and snap frozen on a metal bar cooled by dry ice. Frozen tissue blocks were then sectioned at 10 μm on a Shandon Cryotome E (Thermo Fisher Scientific). Sections were stored at −80°C until use.

Frozen slides were allowed to thaw at room temperature for 10 min before proceeding with staining. Sections were blocked in PBS containing 3% (v/v) normal goat serum (0.1% (v/v) Triton X-100 in PBS for 30 mins at RT. Sections were incubated overnight at 4°C with anti-CD3 (17A2), -B220 (RA3-6B2) (both from BD Biosciences), and/or -F4/80 (Thermo Fisher Scientific) antibodies diluted in 3% (v/v) normal goat serum (Cell Signaling) in PBS. Sections were washed 3 times in PBS, and the incubated with streptavidin AF555 (Thermo Fisher Scientific) for 1 hour at RT. Sections were washed 3 times in PBS and then mounted using Prolong Gold (Thermo Fisher Scientific). Sections were imaged on a Deltavision Personal (GE Healthcare Life Sciences, Taipei, Taiwan) and image analysis was performed using FIJI software v1.51w (NIH Research Services, Bethesda, MD, USA).

### Determination of the intrasplenic localisation of transferred T cells and enumeration of hepatic granulomas

The number of adoptively transferred TCR Tg CD4^+^ T cells, defined based on their dual expression of GFP and CD3, were counted using the cell counter plugin on FIJI software v1.51w. Two non-consecutive sections with similar size from each spleen were analyzed. The number of double positive T cells was quantified in 3 different distinct regions of the spleen: T cell zone (CD3^+^), B cell follicles (CD3^−^F4/80^−^) and the red pulp (F4/80^+^). The number of hepatic granulomas (CD3^+^DAPI^+^F480^+^ aggregates with a size greater than 50 μM) were counted in ten non-consecutive sections. Section selected for counting were more than 50 μM apart to avoid double counting of the same hepatic granulomas.

### Proliferation Modeling

Quantification of cell numbers for *in vitro* was performed by adding a known number of Rainbow Beads (BD) prior to sample acquisition. FlowJo 10 was used to model cell division numbers respective to an unstimulated control. The same gating of cell division numbers were used for responding samples within the time-point. This was used to determine the proportion of cells in each division and at each time-point. 1 × 10^4^ Rainbow Beads were included prior to flow cytometric acquisition to enumerate the number of cells in each division. Since only a proportion of a starting population of cells will have participated in cell division by the end of an experiment, the precursor cohort method allows us to trace cells that have actually divided (precursor cohort) to quantify the time required to reach first division and the rate of subsequent cell cycles. The methods are described by (8) and described here in brief. Precursor cohort numbers (C_i_) are quantified in accordance to the equation below.

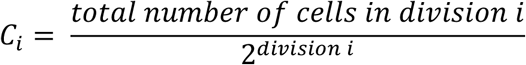

The number of precursor cells in each division at each time point can be plotted. The average division number of cells is estimated by using the equation below. Here, K is the maximum number of divisions that can be resolved.

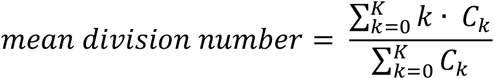

The mean division number is plotted against time. When a trend line is fitted to the data using linear regression analysis, the equation of the line (*y* = *mx*) can be used to calculate the time to first division and the rate of subsequent divisions. Here, *y* is the division number, *m* the gradient, and *x* is time. Using this equation, we can work out the time (x) to first division by substituting y for division number 1. Subsequent division times are calculated by taking the inverse of the gradient:

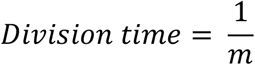

### Estimating the mean distribution of times to first division and T-bet expression

The proportion of undivided or T-bet-negative cells across different time points is fitted with an inverted cumulative lognormal distribution. The best-fit values of the mean (μ) and standard deviation (sd.) for this distribution were used to determine the probability lognormal distribution of times to first division (8, 23, 24) or T-bet expression. The μ of the probability distribution provides an estimate for the time to first division or T-bet expression.

### Statistical Analysis

All statistical analyses were performed using Prism 7 (GraphPad Software, San Diego, CA, USA). Statistical differences between two groups were evaluated using the Student’s t test. Differences between more than two groups were calculated using 2way analysis of variance (ANOVA) with Tukey’s multiple comparisons test. Results with p < 0.05 were deemed statistically significant, where; * p < 0.05, ** p < 0.01, *** p <0.001.

## Acknowledgments

We thank E. Pamer and M. Glickman (Memorial Sloan-Kettering Cancer Center) for providing the transgenic T cell mouse lines. We also thank P. Hodgkin and S. Heinzel (Walter and Eliza Hall Institute) and D. Jankovic (NIH) for reading this manuscript. We acknowledge M. Coleman, S. Warner and H. Rathbone for their thoughtful discussion, and M. Soud for proofreading this manuscript. Finally, we are grateful to the Centenary Institute Animal and Flow Cytometry Core facilities for their support.

---

**Figure. S1.**
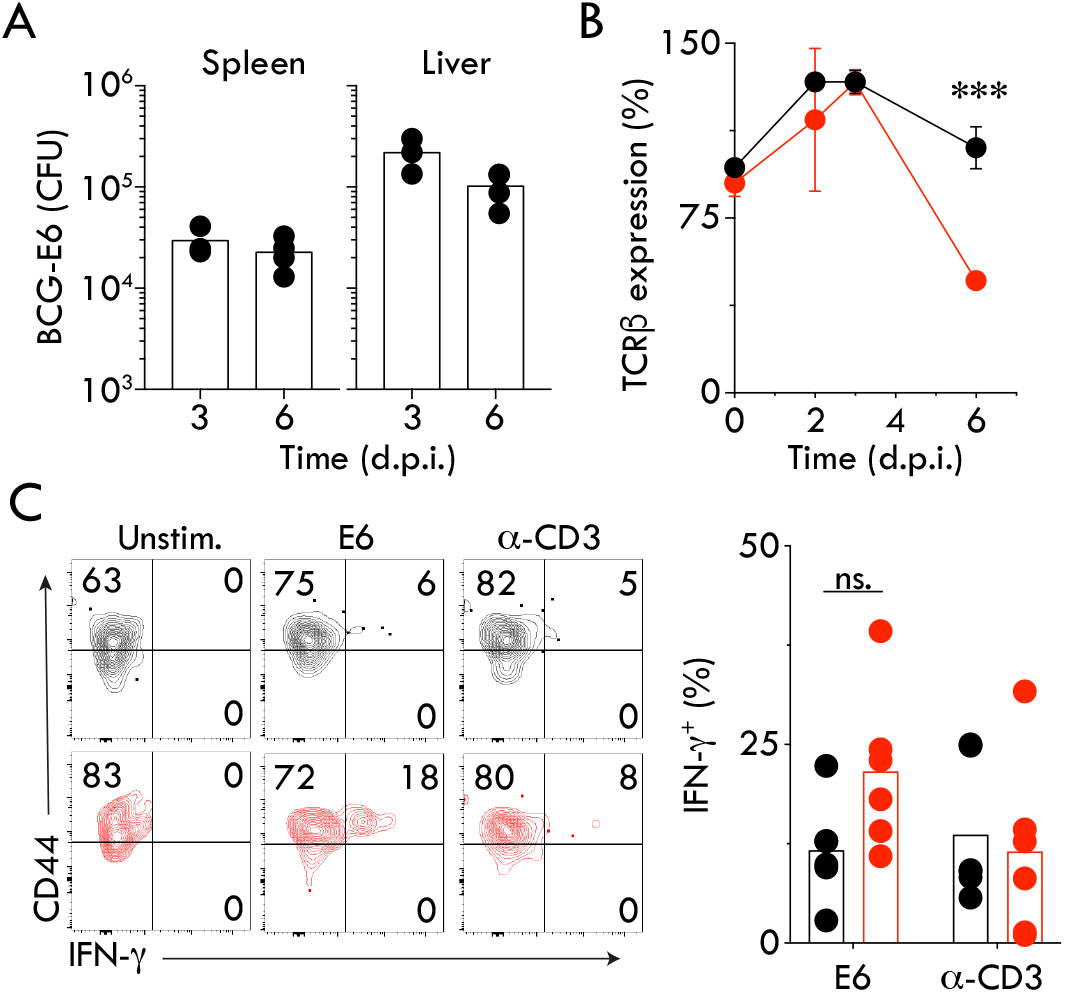
TCR downregulation in high affinity C24 cells does not affect antigen reactivity *ex vivo* and the distribution of BCG-E6 following i.v infection. **(A)** Bacterial loads in the spleen and liver of B6 mice (without the transfer of TCR Tg CD4^+^ T cells) assessed at day 3 and 6 after i.v. BCG-E6 inoculation (n = 4 mice / time-point). **(B-C)** C7 and C24 CD4^+^ cells were transferred i.v. into naive B6 recipients 1 day prior to i.v. inoculation with BCG-E6. **(B)** Flow cytometric analysis of TCRβ expression on C7 and C24 cells in the spleens of mice at the indicated time-points. TCRβ expression on TCR Tg cells is represented as a percentage relative to the MFI of TCRβ on endogenous CD44^−^CD4^+^ T cells. Data are representative of three independent experiments. Symbols denote group means and bars show sem (n = 3 mice / group). **(C)** CD44 and IFN-γ expression in C7 and C24 cells from the spleens of mice 6 days after i.v. inoculation with BCG-E6. Splenocytes were left unstimulated or re-stimulated for 5 h with either the E6 peptide (1 μg/mL) or soluble agonistic anti-CD3 mAb (1 μg/mL) in the presence of BFA. Representative FACS plots and summary graphs of pooled data from two experiments are shown (n = 6 mice / group). Symbols denote individual mice and the open bars represent group means. Black and red flow cytometry plots or symbols represent C7 and C24 cells, respectively. Statistical differences between C7 and C24 cells were determined using the Student’s t test (p*** < 0.001).

**Figure. S2.**
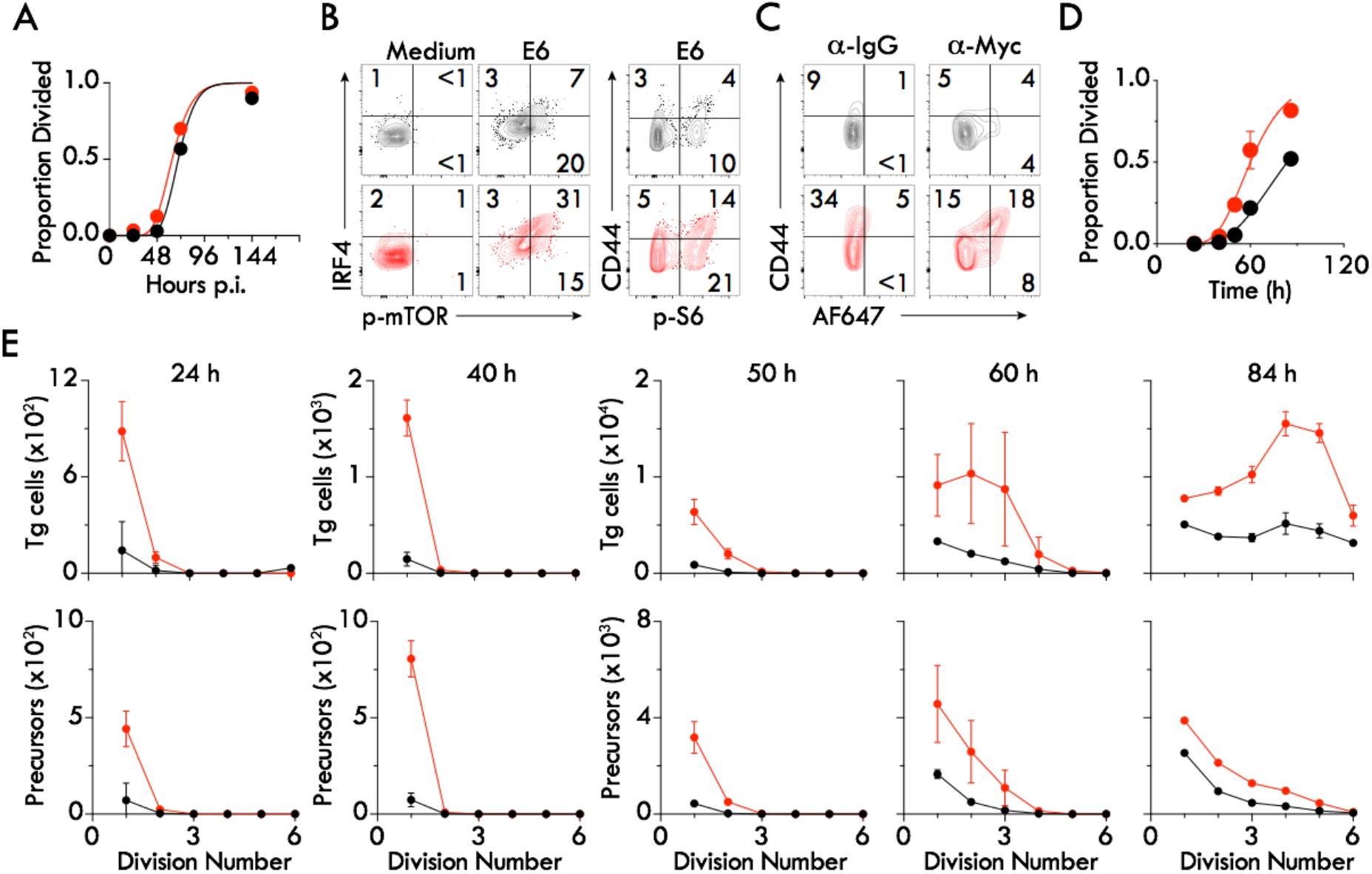
T cell activation and modelling of cell division kinetics. **(A)** C7 and C24 CD4^+^ cells were transferred i.v. into naive B6 recipients 1 day prior to i.v. inoculation with BCG-E6. At each time-point, the average proportion of undivided C7 and C24 cells in the dLN, liver and spleen were determined using flow cytometry, and fitted with an inverted cumulative lognormal distribution (C7: μ = 4.243, sd. = 0.1989; C24: μ = 4.148, sd. = 0.2441) and used to estimate the best fit log normal distribution describing the time to first division (Fig. 2D). Lines and symbols that are black or red represent C7 and C24 cells, respectively. Data shown are representative of two independent experiments with similar results. Symbols denote the group mean (n = 3 - 4 mice / group). **(B-E)** CTV-labelled C7 and C24 CD4^+^ T cells were cultured with CD4-depleted splenocytes in the presence of IL-12, and anti-IL-4 and anti-IFN-γ mAbs, with or without E6 peptide. **(B)** Flow cytometric analysis of IRF4 and phosphorylated mTOR (p-mTOR), CD44 and phosphorylated ribosomal S6 (p-S6) at 24 h. **(C)** Expression of CD44 and Myc 24 h after peptide activation was assessed by incubating C7 and C24 cells with either anti-Myc or rabbit IgG istotype control primary antibodies prior to staining with anti-rabbit IgG conjugated to AF647. Flow cytometry plots that are black or red represent C7 and C24 cells, respectively. Data shown are representative of two independent experiments with similar results. **(D)** The proportion of undivided C7 and C24 cells at each time point was calculated and was fitted with an inverted cumulative lognormal distribution (C7: μ = 4.427, sd. = 0.3736; C24: μ = 4.09, sd. = 0.3013) and used to estimate the best fit log normal distribution describing the time to first division. Lines and symbols that are black or red represent C7 and C24 cells, respectively. Data shown are representative of two independent experiments with similar results. Symbols denote the mean and bars represent the sd of triplicate cultures. **(E)** The number of C7 and C24 cells used to calculate their respective precursor cohorts in each division at the indicated time points was determined by flow cytometric analysis of CTV profiles and counting beads. The numbers in division 0 were excluded to better highlight time-dependent progression through division numbers. **(D-E)** Data shown are representative of three independent experiments with similar results. Lines and symbols that are black or red represent C7 and C24 cells, respectively. Symbols denote the mean and the error bars show the sd. of triplicate cultures.

**Figure. S3.**
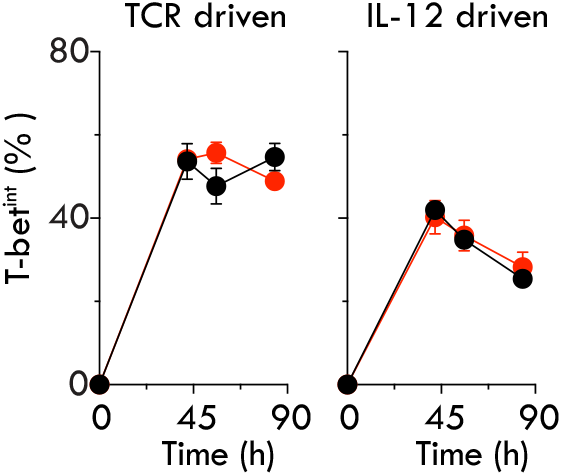
Kinetic analysis of intermediate T-bet-expressing T cell populations. Purified C7 and C24 CD4^+^ T cells were cultured with CD4-depleted splenocytes in the presence or absence of E6 peptide (0.05 μg/mL), and exposed to TCR-driven (anti-IL-12p70, anti-IL-4 and anti-IFN-γ mAbs) or IL-12-driven (5 ng/mL IL-12p70 and anti-IFN-γ and anti-IL-4 mAbs) culture conditions. C7 and C24 populations expressing intermediate levels of T-bet (T-bet^int^) was determined by flow cytometry at the indicated time points. Data are representative of two independent experiments with similar results. Lines and symbols that are black or red represent C7 and C24 cells, respectively. Symbols denote the mean and the error bars show the sd. of triplicate cultures.

**Figure. S4.**
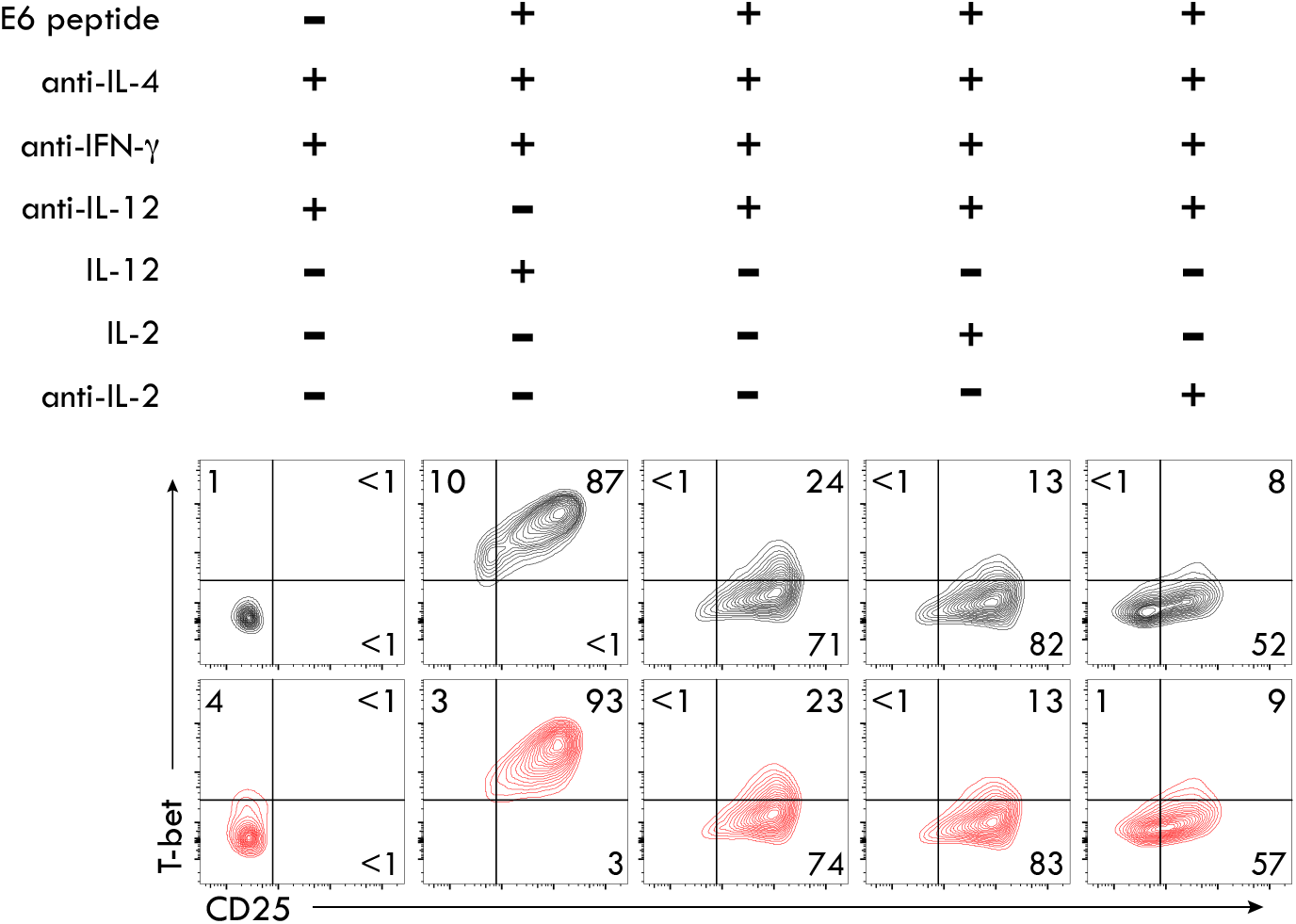
IL-2 minimally affects T-bet expression *in vitro*. Purified C7 and C24 CD4^+^ T cells were cultured with CD4-depleted splenocytes in the presence or absence of E6 peptide (0.05 μg/mL). Where indicated the T cells were incubated anti-IL-12p70, anti-IL-4, anti-IL-2 and anti-IFN-γ mAbs, IL-12 and IL-2. T-bet and CD25 expression was analysed at 72 h following stimulation. Flow cytometry plots that are black and red represent C7 and C24 cells, respectively. Data are representative of two independent experiments with similar results.

